# mRNA vaccine CVnCoV protects non-human primates from SARS-CoV-2 challenge infection

**DOI:** 10.1101/2020.12.23.424138

**Authors:** Susanne Rauch, Karen Gooch, Yper Hall, Francisco J. Salguero, Mike J. Dennis, Fergus V. Gleeson, Debbie Harris, Catherine Ho, Holly E. Humphries, Stephanie Longet, Didier Ngabo, Jemma Paterson, Emma L. Rayner, Kathryn A. Ryan, Sally Sharpe, Robert J. Watson, Stefan O. Mueller, Benjamin Petsch, Miles W. Carroll

## Abstract

The ongoing severe acute respiratory syndrome coronavirus-2 (SARS-CoV-2) pandemic necessitates the fast development of vaccines to meet a worldwide need. mRNA-based vaccines are the most promising technology for rapid and safe SARS-CoV-2 vaccine development and production. We have designed CVnCoV, a lipid-nanoparticle (LNP) encapsulated, sequence optimised mRNA-based SARS-CoV-2 vaccine that encodes for full length, pre-fusion stabilised Spike protein. Unlike other mRNA-based approaches, CVnCoV exclusively consists of non-chemically modified nucleotides and can be applied at comparatively low doses. Here we demonstrate that CVnCoV induces robust humoral and cellular responses in non-human primates (NHPs). Animals vaccinated with 8 μg of CVnCoV were protected from challenge infection with SARS-CoV-2. Comprehensive analyses of pathological changes in challenged animals via lung histopathology and Computed Tomography (CT) scans gave no indication of enhanced disease upon CVnCoV vaccination. These results demonstrate safety, immunogenicity, and protective efficacy of CVnCoV in NHPs that extend our previously published preclinical data and provide strong support for further clinical testing in ongoing phase 2b/3 efficacy studies.

## Introduction

One year after the first cases of severe acute respiratory syndrome coronavirus 2 (SARS-CoV-2) were identified, a pandemic of >70 million confirmed infections has spread worldwide ^1^. Coronavirus infectious disease 2019 (COVID-19) has so far resulted in 1.7 million deaths ^1^, highlighting the urgent need for rapid vaccine development. Fifty-six vaccine candidates are currently in clinical development ^2^, with the first highly promising clinical efficacy data available ^3,4^. mRNA vaccine technologies allow both rapid development and large-scale production ^5^ and have recently emerged as the most promising technology to tackle this worldwide crisis ^6 7 8 9, 10^.

CureVac’s mRNA technology termed RNActive® is based on non-chemically modified, sequence engineered mRNA, and is designed to allow rapid development of human vaccines to emerging diseases. RNActive® vaccines formulated with protamine have been employed in the development of cancer immunotherapy ^11 12 13 14^ as well as in the context of prophylactic vaccines ^15 16 17 18^. A first-in-human phase 1 study using this formulation in a rabies mRNA vaccine demonstrated proof of concept of chemically unmodified mRNA vaccines in humans and found significant levels of rabies neutralizing antibodies ^19^. Further studies established significant improvements of vaccine efficacy by encapsulating the mRNA in lipid nanoparticles (LNPs). These improvements were demonstrated in preclinical models ^16^ and a phase 1 clinical trial, where two 1 or 2 μg doses elicited immune responses comparable to a three-dose regimen of a licensed rabies vaccine ^20^.

RNActive® technology has now been applied to the development of CVnCoV; an LNP formulated mRNA-based vaccine against SARS-CoV-2. The mRNA component of the vaccine consists of non-chemically modified nucleotides and has been optimised for high expression of the encoded protein and moderate activation of innate immune responses. CVnCoV encodes for full-length Spike (S) protein with two proline mutations (S-2P) aimed to stabilise the protein in the pre-fusion conformation, as previously described for MERS- and SARS-CoV-1 ^21 22^. This protein, a trimeric glycoprotein on the viral surface, is responsible for receptor binding and viral entry ^23 24, 25^ and represents a critical target for viral neutralizing antibodies ^26 27^.

Preclinical results have shown the induction of robust cellular and humoral immune responses in CVnCoV vaccinated rodents, as well as the ability to protect against challenge infection in a hamster model ^5^. CVnCoV testing in phase 1 clinical trials demonstrated safety and tolerability as well as full seroconversion in the 12 μg vaccine group two weeks post injection of the second dose. Importantly, this group exhibited comparable virus neutralizing antibody titres to the median titres observed in convalescent sera from COVID-19 patients exhibiting multiple symptoms ^28^. These results supported further clinical testing of the vaccine currently in phase 2b/3 studies that will investigate the efficacy, safety, and immunogenicity of the candidate vaccine CVnCoV.

Here, we evaluate CVnCoV efficacy in a rhesus macaque SARS-CoV-2 challenge model. Non-human primates develop mild clinical disease with high levels of viral replication in both the upper and lower respiratory tract and pathological changes indicative of viral pneumonia upon infection with SARS-CoV-2 ^29 30^. In this study we show that CVnCoV had protective impact against challenge with 5 x 10^6^ PFU via the IN and IT routes in an NHP *in vivo* model of COVID-19. Protective endpoints include significantly reduced virus load and protection against lung pathology.

## Results

### Immunogenicity

Eighteen rhesus macaques of Indian origin were divided into three groups of six, each comprising three males and three females. Animals were vaccinated twice with either 0.5 μg or 8 μg LNP-formulated mRNA encoding SARS-CoV-2 S-2P (CVnCoV) or remained unvaccinated prior to challenge ^30^ with wild type SARS-CoV-2 (Victoria/1/2020) four weeks after the second vaccination (Figure 1A).

**Figure 1:**
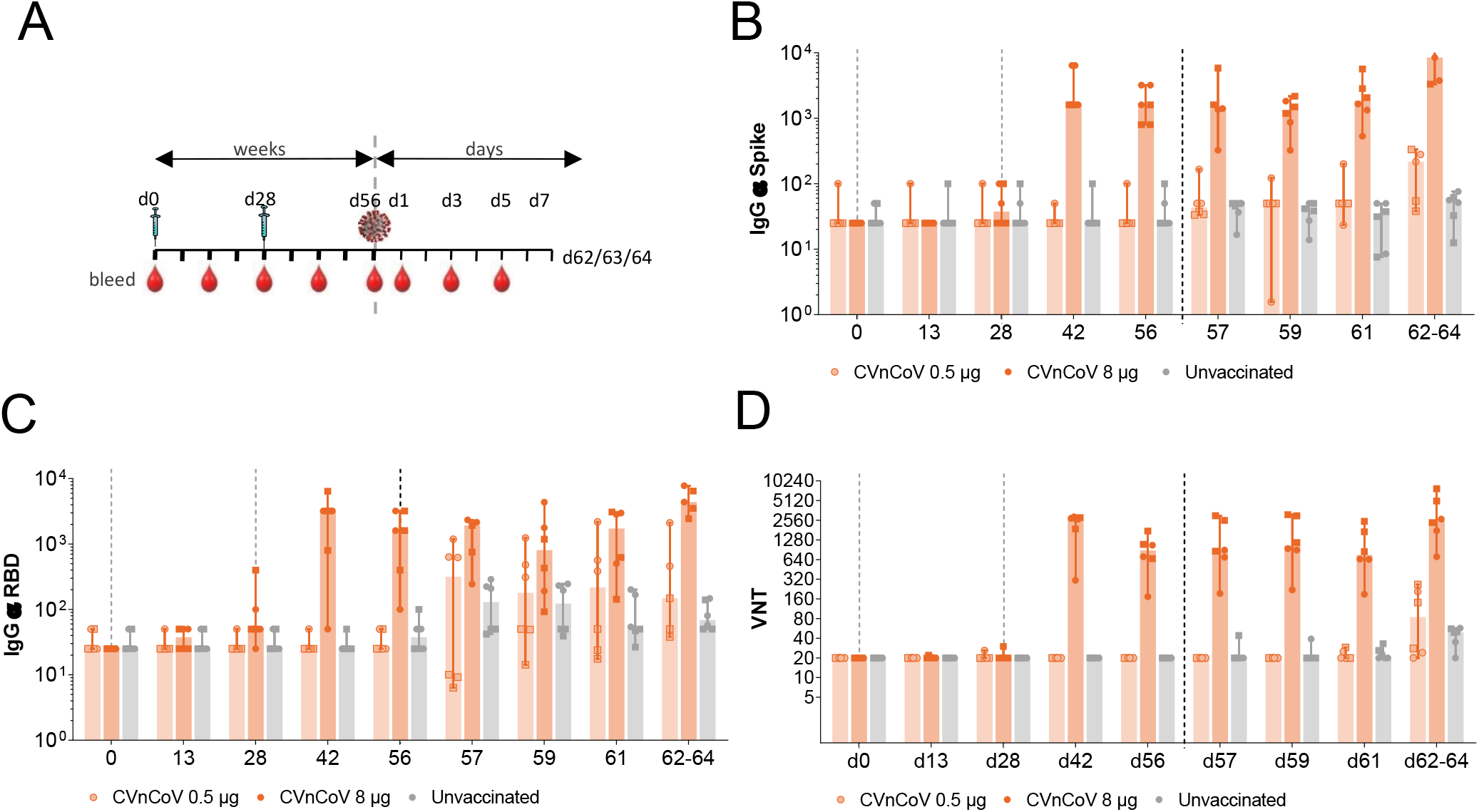
CVnCoV induces humoral response in non-human primates. (A) Schematic drawing of study setup. Rhesus macaques (n=6; 3 male, 3 female/group) were vaccinated IM on day 0 and day 28 with 0.5 μg or 8 μg of CVnCoV or remained unvaccinated. All animals were challenge with 5.0 x 106 PFU of SARS-CoV-2 on d56. Two animals of each group were terminated on d62, d63 and d64, respectively (B) Trimeric Spike protein or (C) RBD specific binding IgG antibodies, displayed as endpoint titres at different time points as indicated (C) Virus neutralising antibodies determined via focus reduction neutralisation test at different time points as indicated. All values are displayed as median with range. Square symbols represent male, round symbols female animals. Dotted lines represent vaccinations and challenge infection, respectively. RBD receptor binding domain; VNT virus neutralising titre

Analysis of binding titres to either a trimeric form of the S protein or the isolated receptor binding domain (RBD) showed a measurable increase in spike (Figure 1B) and RBD-specific IgG titres (Figure 1C) in animals vaccinated with 8 μg after a single vaccination. A significant increase was observed in IgG titres after the second vaccination on study day 42, with animals exhibiting median endpoint titres of 1.6 x 10^3^ and 3.2 x 10^3^ for S and RBD reactive antibodies, respectively (Figures 1 B and C). A further increase of spike- and RBD specific IgG titres was seen upon challenge in this group, particularly in serum collected at the time of termination (study days 62, 63, and 64).

As expected, no significant increase in spike or RBD-specific IgG antibodies was seen in the 0.5 μg CVnCoV (intentionally sub-optimal) dose or in the unvaccinated control group during the vaccination phase. However, a gradual increase in Spike and RBD specific IgG titres was observed at each of the sampling points in animals vaccinated with 0.5 μg CVnCoV after challenge (Figures 1 B and C). No increase in Spike-or RBD specific IgG titres was observed in the unvaccinated controls (Figure 1B and C).

In agreement with the induction of binding antibodies, robust levels of virus neutralising titres (VNTs) were detectable after the second vaccination in the 8 μg group (Figure 1D). VNTs peaked on d42 at median titres of 2.7 x 10^4^. Neutralising antibody titres remained relatively unchanged upon challenge until day 62, 63, and 64 of the experiment. Animals in the 0.5 μg and unvaccinated control groups remained negative before challenge, while SARS-CoV-2 infection induced small increases in antibody titres in 4/6 and 5/6 animals in the 0.5 μg and unvaccinated group, respectively.

In order to assess CVnCoV induced cellular responses, peripheral blood mononuclear cells (PBMCs) isolated at different time points post vaccination and challenge were stimulated with pools of peptides spanning the SARS-CoV-2 spike protein. IFN-γ release of stimulated cells was measured by ELISpot. Analysis of responses to summed pools in the vaccination phase showed increases in spike-specific IFN-γ in 8 μg CVnCoV vaccinated animals, two weeks after the first and, more pronounced, two weeks after the second vaccination (Figure 2A panel 1). Stimulation with ten individual pools each covering approximately 140 amino acids of the S protein demonstrated the induction of cells reactive to peptides across the whole length of S upon vaccination with 8 μg of CVnCoV (Figure 2A panel 3).

**Figure 2:**
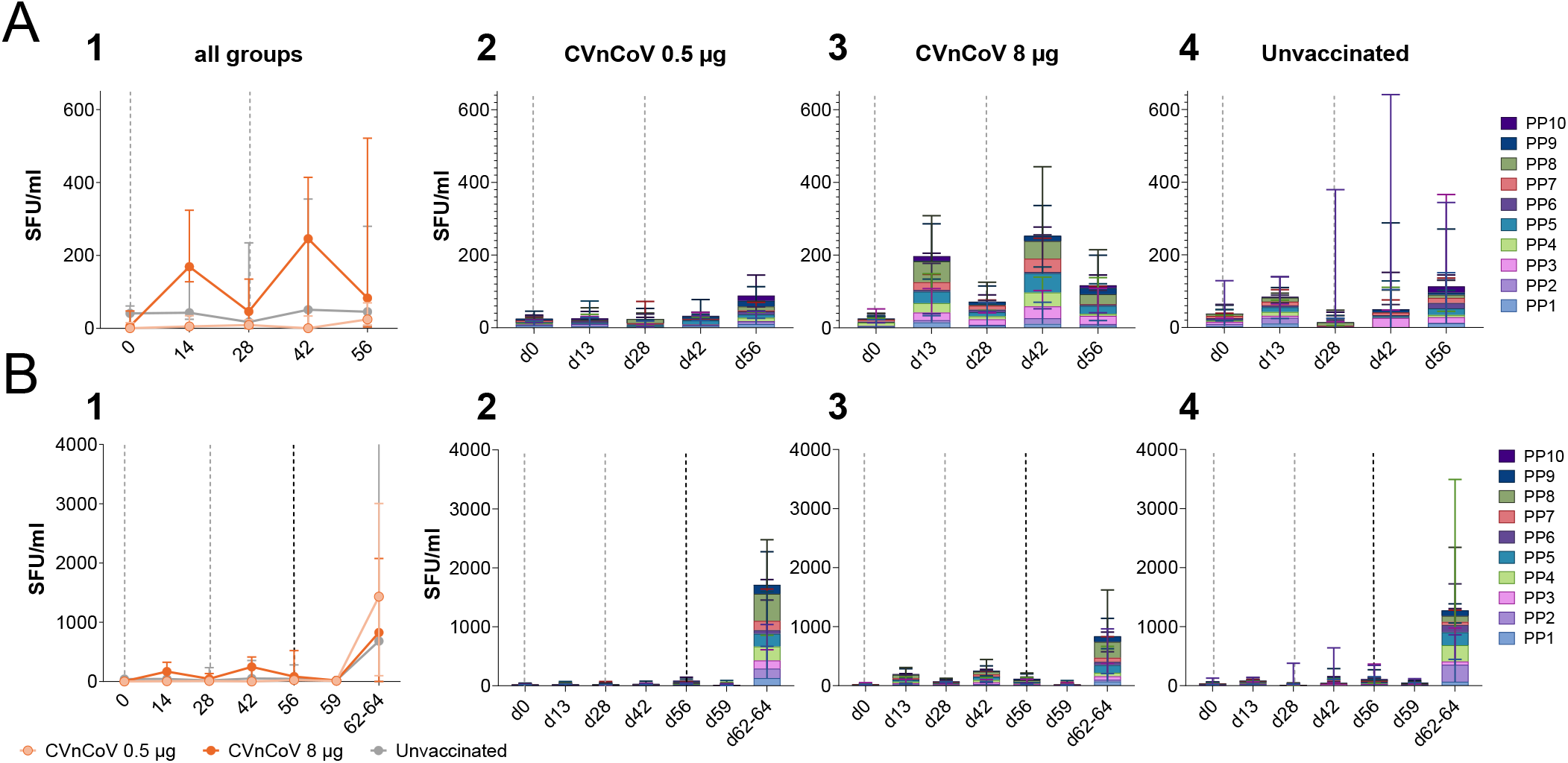
CVnCoV induces cellular responses in non-human primates. PBMCs from 0.5 μg or 8 μg CVnCoV vaccinated or from untreated animals isolated at different time points were re-stimulated with S specific peptide pools ex vivo followed by IFNg ELISpot analysis. (A) IFNg ELISpot before challenge infection on d56. Panel 1 represent results of stimulation with three megapools and shows the summed response covering the whole S protein, panels 2-4 depict stimulation results of ten individual pools covering the entire S protein in each group. (B) IFNg ELISpot until termination on d62-d64. Panel 1 represent results of stimulation with three megapools and shows the summed response covering the whole S protein, panels 2-4 depict stimulation results of ten individual pools covering the entire S protein in each group. SFU spot forming unit; PP peptide pool

There were no clear responses in 0.5 μg CVnCoV or unvaccinated animals in the vaccination phase (Figure 2A panels 1, 2 and 4). One of the female animals in the negative control group showed particularly high IFN-γ secretion after stimulation with peptide pool 2 (covering part of the N-terminal domain (NTD)) throughout the experiment and peptide pool 3 (covering part of the NTD and RBD) on d56 (Figure 2A panel 1 and 4).

Increased spike-specific IFN-γ responses were detectable in all animals on d62-64 post challenge (Figure 2B panels 1-4). Of note, increases of cellular responses in animals vaccinated with 8 μg of CVnCoV were less pronounced than in the other groups, likely indicative of lower levels of viral replication in these animals (Figure 2B panel 3).

### Protective efficacy

Presence or SARS-CoV-2 total RNA in the upper and lower respiratory tract post-challenge was monitored via qRT-PCR (Figure 3). Viral replication in the upper respiratory tract peaked on d59 in unvaccinated animals, which reached median values of 2.7 x 10^7^ cp/ml in nasal swabs (Figure 3A), and remained detectable until termination on d62-d64. No significant difference between viral replication in animals vaccinated with 0.5 μg CVnCoV and unvaccinated control animals was measured in nose swabs. Overall, 8 μg CVnCoV vaccination induced the lowest number of viral RNA copies in the upper respiratory tract, where median values of 2.9 x 10^6^ cp/ml in nasal swabs, respectively, were detectable on d59. However, the difference between the study groups was not statistically significant. Comparable results were generated in throat swabs (Supplementary Figure 1A).

**Figure 3:**
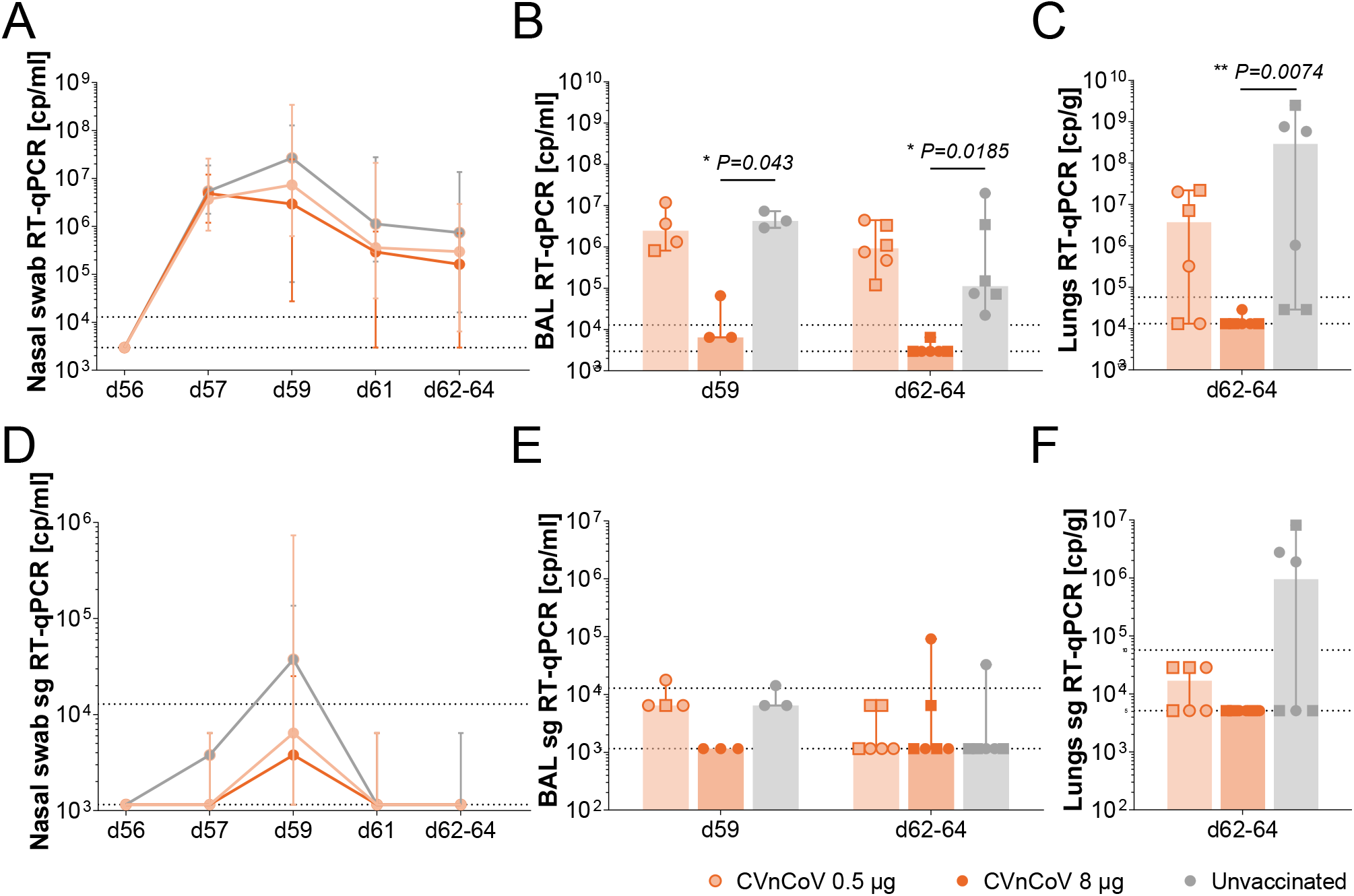
CVnCoV protects non-human primates from challenge infection. (A) Nasal swabs taken at different time points post challenge (B) in life BAL samples taken on d59 and at termination on d62-64 and (C) lung tissue homogenates from d62-64 were analysed for copies of total viral RNA via RT-qPCR. (D) Nasal swabs taken at different time points past challenge (E) in life BAL samples taken on d59 and at termination on d62-64 and (F) lung tissue homogenates from d62-64 were analysed for copies of subgenomic viral RNA via RT-qPCR. Values are depicted as medians with range. Square symbols represent male, round symbols female animals. Lower and upper dotted lines represent LLOD and LLOQ, respectively. Kruskall-Wallis ANOVA followed by Dunn’s test was used to compare groups and P values are shown. LLOD lower limit of detection, LLOQ lower limit of quantification, RT-qPCR Reverse transcription-quantitative polymerase chain reaction

Additional analyses assessing subgenomic (sg) RNA via qRT PCR indicative of viral replication yielded overall low sgRNA levels in the upper respiratory tract. Values peaked on d59 and returned to baseline on d62 in all animals. In nasal swabs, sg RNA levels were lowest in CVnCoV vaccinated animals with values of 0.4 x 10^4^ compared to 3.7 x 10^4^ cp/ml in unvaccinated control animals. Three of 6 animals in the 8 μg CVnCoV vaccinated group remained negative at all time points while 5/6 animals in the unvaccinated group had detectable levels of subgenomic viral RNA (Figure 3D). Analyses of throat swabs showed no significant difference of subgenomic RNA between the groups and median values remained below the lower limit of quantification in all animals (Supplementary Figure 1 B).

Parallel analyses of the lower respiratory tract of in-life (d59) and post-mortem (d62-d64) bronchoalveolar lavage (BAL) samples showed significantly reduced levels of total viral RNA upon 8 μg CVnCoV vaccination at both time points (Figure 3B). Median values of total RNA on d59 and d62-d64 were 4.3 x 10^6^ and 1.1 x 10^5^ cp/ml in the control group, while animals vaccinated with 8 μg of CVnCoV featured median titres of 0.6 x 10^4^ and 0.3 x 10^4^, respectively. RNA levels in BAL were below the lower limit of quantification for all but one animal in the 8 μg CVnCoV group on d59, which featured low RNA counts. Total viral RNA levels in 0.5 μg CVnCoV vaccinated animals were comparable to the control group. Of note, BAL analyses on d59 only depict female animals and one male animal of the unvaccinated group. The remaining animals were excluded from this analysis since suboptimal BAL sampling conditions prevented further evaluation.

The analysis of lung tissue collected at necropsy confirmed results gained in BAL samples. Median titres of 2.9 x 10^8^ cp/g were detectable in the unvaccinated group while all animals in the CVnCoV 8 μg vaccinated groups remained below the lower limit of quantification (Figure 3C). There was no statistically significant difference between animals in the 0.5 μg CVnCoV and the unvaccinated group.

Subgenomic viral RNA analysis in BAL and lung tissue samples yielded comparable results: RNA indicative of replicating virus was detectable in BAL and lung samples of unvaccinated and 0.5 μg CVnCoV vaccinated animals on d59 and d62-d64, respectively. All animals in the 8 μg CVnCoV group were negative in these analyses (Figure 3E and F).

Evaluation of further tissue samples collected at necropsy revealed low but detectable signals of SARS-CoV-2 total RNA in trachea and tonsils of 0.5 μg CVnCoV and unvaccinated animals, while 8 μg CVnCoV vaccinated animals remained negative (Supplementary Figure 1 C and D). No viral RNA was detectable in spleen, duodenum, colon, liver, or kidney in any group (Supplementary Figure 1 E-I).

### Histopathology and imaging

Histopathological analyses of lung samples taken at necropsy showed lesions consistent with infection with SARS-CoV-2 in the lungs of challenged animals (Figure 4). Briefly, the lung parenchyma showed multifocal to coalescing areas of pneumonia surrounded by unaffected parenchyma. Alveolar damage, with necrosis of pneumocytes was a prominent feature in the affected areas. The alveolar spaces within these areas were often thickened. Damaged alveolar walls contained mixed inflammatory cells including macrophages, lymphocytes, viable and degenerated neutrophils, and occasional eosinophils. Alveolar oedema and alveolar type II pneumocyte hyperplasia were also observed. In distal bronchioles and bronchiolo-alveolar junctions, degeneration and sloughing of epithelial cells were present. In the respiratory epithelium of larger airways, occasional focal, epithelial degeneration, and sloughing were observed. Low numbers of mixed inflammatory cells, comprising neutrophils, lymphoid cells, and occasional eosinophils, infiltrated bronchial and bronchiolar walls. In the lumen of some airways, mucus admixed with degenerated cells, mainly neutrophils and epithelial cells, was seen. Within the parenchyma, perivascular and peribronchiolar cuffing were also observed, with mostly lymphoid cells comprising the infiltrates. No remarkable changes were observed in non-pulmonary tissues.

**Figure 4:**
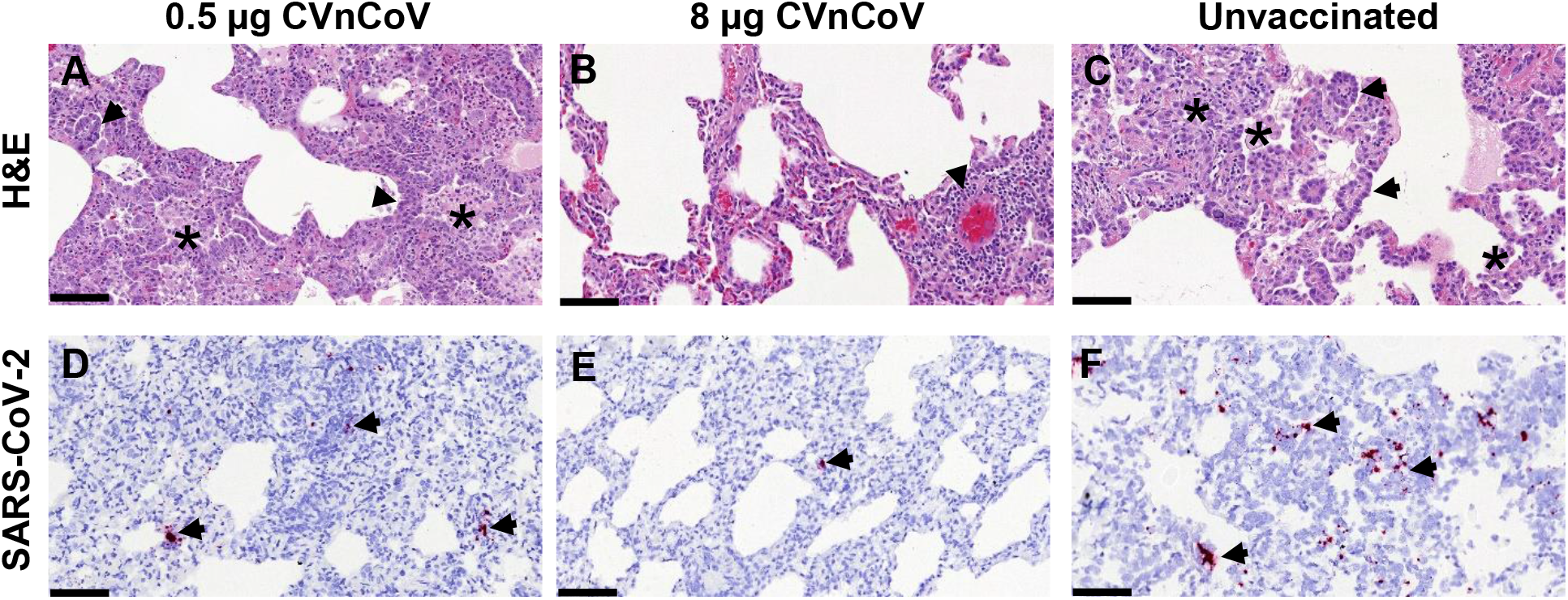
Exemplary sections showing histopathology (H&E) and SARS-CoV-2 *in situ* hybridisation (ISH). A. Alveolar necrosis and inflammatory exudates (*) in the alveolar spaces and type II pneumocyte hyperplasia (arrows). B) Mild perivascular cuffing (arrow). C) Inflammatory cell infiltration in the alveolar spaces and the interalveolar septa (*) and type II pneumocyte hyperplasia (arrows). D) SARS-CoV-2 ISH staining in abundant cell within inflammatory foci (arrows). E. SARS-CoV-2 ISH staining in a single cell within an interalveolar septum (arrow). F. Abundant foci of SARS-CoV-2 ISH stained cells within the alveolar lining and the interalveolar septa (arrows). Bar = 100μm. ISH *in situ* hybridisation

In agreement with reduced levels of viral RNA, the evaluation of lung samples using a histopathology scoring system showed a significant reduction in severity of lung lesions in CVnCoV vaccinated animals compared to 0.5 μg CVnCoV vaccinated and unvaccinated groups (Figure 5A and B).

**Figure 5:**
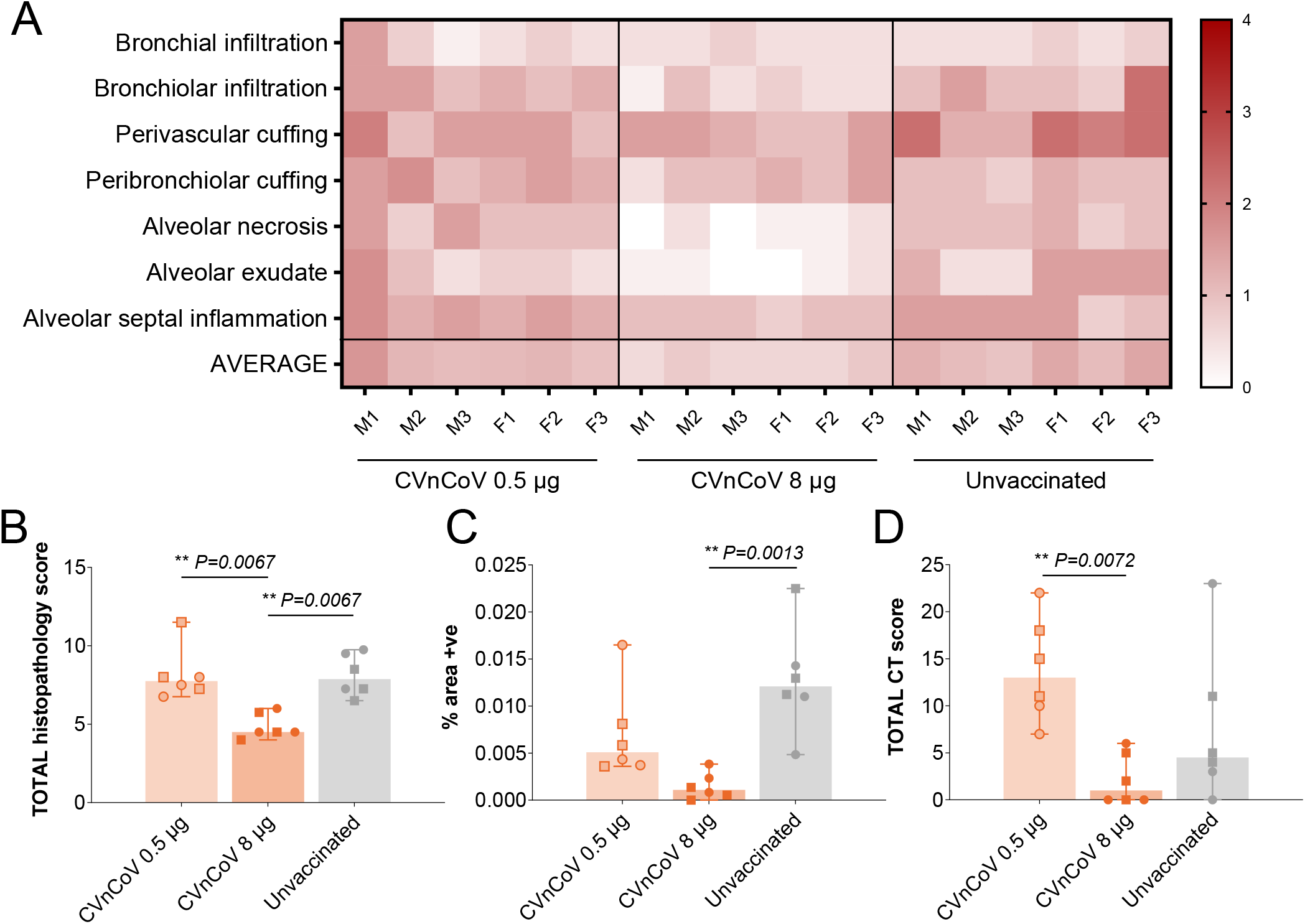
Vaccination with 8 μg of CVnCoV protects the lungs from pathological changes upon viral challenge. (A) Heat map showing scores for each lung pathology parameter and the average score for each animal from all groups as indicated. Severity ranges from 0 to 4: 0=none; 1=minimal; 2=mild; 3=moderate and 4=marked/severe. (B) Graph representing the cumulative score for all the lung histopathology parameters from each animal. (C) Presence of viral RNA in lung tissue sections from all animals expressed as percentage of ISH (RNAScope) positive staining area of lung section. (D) Cumulative score of lung pathology detected via CT radiology. Square symbols represent male, round symbols female animals. Box and whiskers indicate median with range. Kruskall-Wallis ANOVA followed by Dunn’s test was used to compare groups and *P* values are shown. ISH *in situ* hybridisation; M1-3: male animal 1-3; F1-3: female animal 1-3

Viral RNA was observed in alveolar epithelial cells and within the inflammatory cell infiltrates (Figure 4). The quantity of virus RNA observed by *in situ* hybridisation (ISH) was also significantly reduced in 8 μg CVnCoV vaccinated animals when compared to 0.5 μg CVnCoV and unvaccinated (Fig 5C).

To gain in-life view of pathological changes induced upon SARS-CoV-2 infection in the complete lung, CT scanning was performed prior to challenge and post-challenge on study day 61. Overall, the apparent level of disease was relatively mild and only affected less than 25% of the lung. Post-challenge, abnormalities in the lung were detected in 6 of 6 animals in the 0.5 μg CVnCoV group, and 5 of 6 in the unvaccinated control group, while only 3 of 6 animals vaccinated with 8 μg CVnCoV exhibited detectable changes. Lowest levels of total scoring in CT scans were seen in animals vaccinated with 8 μg of CVnCoV (Figure 5D). Of note, highest scores were seen in the 0.5 μg CVnCoV group in this analysis. However, values were not statistically different to the control group.

## Discussion

Our data demonstrate that CVnCoV is safe and highly immunogenic in rhesus macaques. Vaccination with 8 μg CVnCoV elicited robust humoral responses, known to correlate with protection in this model as RBD-specific monoclonal antibodies ^31^ and purified IgGs from convalescent animals ^32^ are able to protect naïve rhesus macaques from challenge infection. Here we show the induction of high levels of S and RBD specific binding and virus neutralising antibodies upon two 8 μg injections of CVnCoV. While a sub-optimal dose of 0.5 μg was unable to elicit detectable levels of antibodies before challenge, infected animals raised specific antibody responses faster than the unvaccinated control group, indicative of a previously induced priming response in these animals. These data extend our preclinical finding in mice and hamsters, that elicited robust, dose-dependent humoral immune responses upon vaccination with CVnCoV using comparable or lower doses of CVnCoV ^5^. In addition, our results support previously published interim phase I clinical data that demonstrated the induction of humoral responses with two doses up to 12 μg ^28^.

Furthermore, data presented here demonstrated the strong induction of S specific cellular responses in CVnCoV vaccinated animals via IFNγ ELISpot. In animals vaccinated with 8 μg of CVnCoV, increasing responses against peptides covering the whole length of the S protein were elicited after first and second vaccination. These data are in line with our previous reports in mice that demonstrated the ability of CVnCoV to induce high S-specific CD4+ and CD8+ T cell responses ^5^. The generation of robust T cell responses is likely to support vaccine efficacy against SARS-CoV-2. Recent data have demonstrated that CD8+ T cells contribute to viral control in a rhesus macaque model ^32^. While the exact role of T cells in the context of COVID-19 is still under some debate, T cell responses to SARS-CoV-2 are readily detectable in humans and may play a role in long term protection ^33 34 35^.

Of note, the analysis of cellular responses revealed one animal in the unvaccinated group that mainly reacted against a single peptide pool of S covering amino acids 129-267 which represent part of the NTD of S. Responses to other immunological read outs, i.e. specific antibodies and nucleocapsid and membrane antigens in ELISpot, were not high pre-challenge, ruling out a prior infection of this particular animal with SARS-CoV-2. While the reason for this observation remains unclear, a possible explanation is cross-reactive T cell epitope of a pre-existing response to a seasonal coronavirus infection.

Quantification of viral RNA copies upon challenge infection demonstrated a reduction of viral replication in the upper respiratory tract in 8 μg CVnCoV vaccinated animals. Importantly, this vaccine dose was able to protect the lungs of challenged animals. Protection was both demonstrated by undetectable levels of viral RNA and by reduced pathological changes upon challenge infection compared to unvaccinated animals. More effective protection of the lower respiratory tract (LRT) compared to the upper respiratory tract (URT) in the presence of a robust immune responses is in line with our previously published results in hamsters ^5^ and with reports of other mRNA-based SARS-CoV-2 vaccines in NHP challenge models ^6 7^.

In terms of vaccine safety, the injection of 0.5 μg or 8 μg of CVnCoV elicited no adverse reactions to vaccination and no differences in weight or temperature were observed between groups during the vaccination phase of the study (data not shown), supporting a favourable safety profile of the vaccine in rhesus macaques at the doses used.

Previous vaccination studies in the context of SARS- and Middle East Respiratory Syndrome (MERS)-CoV suggest an association with vaccine enhanced disease in animal models (reviewed in ^36 37^), highlighting the need for careful evaluation of this potential risk during SARS-CoV-2 vaccine development. However, no signs of vaccine enhanced disease (VED), post challenge, were observed in this study. To date, neither preclinical ^38 6 39 40^, nor clinical evidence indicating a risk of VED in the context of SARS-CoV-2 infections has been reported. Vaccine enhanced disease can be caused by antibodies (antibody-dependent enhanced disease, ADE (reviewed in ^41^) as previously described for a feline coronavirus ^42^. Such antibodies most likely possess non-neutralising activity, and enhance viral entry causing increased viral replication and disease exacerbation. Results presented here give no indication of increased viral replication in animals vaccinated with CVnCoV. Importantly, enhanced replication in the respiratory tract or distal organs such as spleen, duodenum, colon, liver, or kidney was not detectable in the 0.5 μg group of the study. These animals featured low levels of S binding but undetectable levels of VNTs upon challenge infection, creating conditions under which ADE could hypothetically occur.

Another cause of disease enhancement may be vaccine-associated enhanced respiratory disease (VAERD) that is hallmarked by increased inflammation due to Th2-biased immune responses and high ratios of non-neutralising to neutralising antibodies (reviewed in ^43 41 44^). Analysis of lung pathology in CVnCoV vaccinated animals demonstrated protectivity of 8 μg CVnCoV and gave no indication for increased inflammation and pathological changes in sub-optimally dosed animals.

Results presented here extend our knowledge of CVnCoV safety, immunogenicity and protective efficacy in a highly relevant model system for SARS-CoV-2. The overall outcome of the study in non-human primates in terms of immunogenicity, protective efficacy, and pathology are comparable to results in the hamster model, providing support for hamsters as a model system for SARS-CoV. We conclude that CVnCoV is highly efficacious at a low dose of 8 μg in a COVID-19 NHP challenge model while being safe at both doses tested with lack of any indication of disease enhancement. In alignment with the phase I interim safety and immunogenicity data of CVnCoV ^28^, these results provide strong support for further evaluation in the ongoing phase 2b/3 HERALD clinical trial.

## Materials and Methods

### mRNA vaccines

The mRNA vaccine is based on the RNActive® platform (claimed and described in e.g. WO2002098443 and WO2012019780) and is comprised of a 5’ cap structure, a GC-enriched open reading frame (ORF), 3’ UTR, polyA tail. It does not include chemically modified nucleosides. Lipid nanoparticle (LNP)-encapsulation of mRNA was performed by Polymun Scientific Immunbiologische Forschung GmbH (Klosterneuburg, Austria). The LNPs used in this study are particles of ionizable amino lipid, phospholipid, cholesterol and a PEGylated lipid. The mRNA encoded protein is based on the spike glycoprotein of SARS-CoV-2 NCBI Reference Sequence NC_045512.2, GenBank accession number YP_009724390.1 and encodes for full length S featuring K986P and V987P mutations.

### Animals

Rhesus macaques *(Macaca mulatta),* Indian origin, were obtained from a UK Home Office approved colony. In total, this study included 18 animals (9 male, 9 female) with a weight of > 4.5 kg and an age of 3-6 years.

All animal work was conducted at Public health England (PHE) Porton Down in compliance with the UK Home Office Animals (Scientific Procedures) Act 1986 with licensed individuals performing all procedural techniques. The study was conducted under the authority of a UK Home Office approved project licence that had been subject to local ethical review at PHE Porton Down by the Animal Welfare and Ethical Review Body (AWERB). Animals were housed in compatible social groups, in cages in accordance with the UK Home Office Code of Practice for the Housing and Care of Animals Bred, Supplied or Used for Scientific Procedures (2014) and National Committee for Refinement, Reduction and Replacement (NC3Rs) Guidelines on Primate Accommodation, Care and Use, August 2006.

### Vaccinations

Animals were injected intramuscularly (IM) in the bicep muscle of the upper arm with 0.5 μg or 8 μg of LNP-formulated SARS-CoV-2 mRNAs (CVnCoV), in a volume of 0.5 ml, on d0 and d28 of the study or left untreated as negative controls.

### SARS-CoV-2 challenge infection

The challenge agent used in this study was SARS-CoV-2 virus, VERO/hSLAM cell passage 3 (Victoria/1/2020) ^45^, titre 2.4 x 10^7^ PFU/ml, passaged from material (P1) generously provided to PHE by The Doherty Institute, Melbourne, Australia. All animals were challenged with a dose of 5.0 x 10^6^ PFU of SARS-CoV-2 by applying 2 ml of virus preparation to the pre-carinal section of the trachea using a bronchoscope followed by 1 ml applied intranasally (0.5 ml/nostril). Two animals of each group were followed for 6, 7 or 8 days post challenge (p.c.) and euthanised on day 62, 63 or 63 of the experiment.

### IgG ELISA

A full-length trimeric and stabilised version of the SARS-CoV-2 Spike protein was supplied by Lake Pharma (#46328). Recombinant SARS-CoV-2 Receptor-Binding-Domain (319-541) Myc-His was developed and kindly provided by MassBiologics. Recombinant SARS-CoV-2 Spike- and RBD-specific IgG responses were determined by ELISA. High-binding 96-well plates (Nunc Maxisorp, 442404) were coated with 50 μl per well of 2 μg/ml Spike trimer or RBD in 1x PBS (Gibco) and incubated overnight at 4°C. The ELISA plates were washed and blocked with 5% Foetal Bovine Serum (FBS, Sigma, F9665) in 1x PBS/0.1% Tween 20 for 1 hour at room temperature. Serum collected from animals after vaccination had a starting dilution of 1/50 followed by 8, two-fold serial dilutions. Post-challenge samples were inactivated in 0.5% triton and had a starting dilution of 1/100 followed by 8, three-fold serial dilutions. Serial dilutions were performed in 10% FBS in 1x PBS/0.1% Tween 20. After washing the plates, 50 μl/well of each serum dilution was added to the antigen-coated plate in duplicate and incubated for 2 hours at room temperature. Following washing, anti-monkey IgG conjugated to HRP (Invitrogen, PA1-84631) was diluted (1: 10,000) in 10% FBS in 1X PBS/0.1% Tween 20 and 100 μl/well was added to the plate. Plates were then incubated for 1 hour at room temperature. After washing, 1 mg/ml O-Phenylenediamine dihydrochloride solution (Sigma P9187) was prepared and 100 μl per well were added. The development was stopped with 50 μl per well 1M Hydrochloric acid (Fisher Chemical, J/4320/15) and the absorbance at 490 nm was read on a Molecular Devices versamax plate reader using Softmax (version 7.0) All test sample dose response curves were fitted to a 4PL model in Softmax Pro (version 7.0) and the endpoint titre at an OD of 0.5 (defined as reciprocal of the serum dilution required to give an absorbance response of 0.5) was interpolated from each curve. Where results were below the limit of detection, they were assigned a value of 25 for the post immunisation samples and 50 for the post challenge samples. For low samples where the absorbance never reached a value of 0.5, the titre was estimated from the extrapolated portion of the curve. The cut-off was set as the average titre of serum collected from naïve animals (day 0) + 1 Standard Deviation. The cut off was calculated separately for each antigen.

### SARS-CoV-2 focus reduction neutralisation test

Virus neutralising titres were measured in heat-inactivated serum samples (56°C for 30 min). SARS-CoV-2 (Victoria/01/2020, Doherty Institute) at a concentration to give 100 to 250 foci per well in the virus only control wells was mixed 50:50 in 1% FCS MEM with 1 x antibiotic/antimycotic (Gibco, 15240-062) with serum doubling dilutions from 1:20 to 1:640 (or higher dependent on antibody levels) in a 96-well V-bottomed plate. The plate was incubated at 37°C in a humidified box for 1 hour to allow antibodies in the serum sample to bind to the virus. One hundred microlitres of the serum/virus mixture was then transferred to virus susceptible Vero/E6 monolayers in 96-well plates and incubated for a further 1 hour at 37°C in a sealed humidified box. After adsorption, the virus/antibody mixture was removed and 100 μl of 1% w/v CMC in complete media overlay was added. The box was resealed and incubated at 37°C for 24 hours prior to fixing with 100 μl of 20% formalin/PBS solution and fumigation of the plate overnight prior to immunostaining.

Following washing with water using an ELISA washer (BioTek 405 TSUS), residual endogenous peroxidase activity was removed by the addition of 0.3% hydrogen peroxide for 20 min. Plates were then incubated for 1 h with primary/detection SARS-CoV-2 anti-RBD rabbit polyclonal antibody (SinoBiologicals; 40592-T62) diluted 1: 2,000 in PBS. After washing, plates were incubated for 1 h with secondary anti-rabbit HRP-conjugate antibody (Invitrogen; G-21234) diluted 1: 4,000 in PBS. After washing, foci were visualised using TrueBlue™ Peroxidase Substrate (KPL seracare; 5510-0030) after which plates were washed with water and dried. Foci were counted using an ImmunoSpot S6 Ultra-V analyser (CTL) and BioSpot software (7.0.28.4 Professional; CTL) and the results analysed in SoftMax Pro (Molecular Devices; v7.0.3 GxP). Briefly, the count data was expressed as percentage of VOC for each serum dilution, i.e. percentage foci reduction and plotted on a 4-Parameter logistic (4PL) curve. The virus neutralisation titre (VNT) is reported as serum dilution that neutralised 50% of the virus foci.

### ELISpot

Peripheral Blood Mononuclear Cells (PBMCs) were isolated from whole heparinised blood by density gradient centrifugation using Ficoll-Paque Plus (GE Healthcare, USA). An IFN-γ ELISpot assay was used to estimate the frequency and IFN-γ production capacity of SARS-CoV-2-specific T cells in PBMCs using a human/simian IFN-γ kit (MabTech, Nacka, Sweden). The cells were assayed at 2 x 10^5^ cells per well. Cells were stimulated overnight with SARS-CoV-2 peptide pools and ‘megapools’ of the spike protein (Mimotopes, Australia). Peptide sequence was based on GenBank: MN908947.3. Ten peptide pools were used, compriseing of 15 mer peptides, overlapping by 11 amino acids. The three megapools were made up as such: Megapool 1 (MP1) comprised peptide pools 1-3, Megapool 2 (MP2) comprised peptide pools 4-6 and Megapool 3 (MP3) comprised of peptide pools 7-10. All peptides were used at a final concentration of 1.7 μg/ml per peptide. Phorbol 12-myristate (Sigma-Aldrich Dorset, UK) (100 ng/ml) and ionomycin (CN Biosciences, Nottingham, UK) (1 mg/ml) were used as a positive control. Results were calculated to report as spot forming units (SFU) per million cells. All SARS-CoV-2 peptides and megapools were assayed in duplicate and media only wells subtracted to give the antigen-specific SFU. ELISpot plates were analysed using the CTL scanner and software (CTL, Germany) and further analysis carried out using GraphPad Prism (version 8.0.1) (GraphPad Software, USA).

### Bronchioalveolar lavage (BAL)

In-life BAL washes were performed using 10 ml PBS using a bronchioscope inserted to the right side of the lung above the second bifurcation. BAL washes performed post-mortem were conducted on the right lung lobes, after ligation of the left primary bronchus using 20 ml PBS.

### Quantitative Polymerase Chain Reaction

RNA was isolated from nasal swab, throat swabs, EDTA treated whole blood, BAL and tissue samples (spleen, kidney, liver, colon, duodenum, tonsil, trachea and lung). Tissue samples in RNAprotect (Qiagen), were homogenised in a Precellys 24 homogeniser with CK28 Hard tissue homogenizing 2.0 ml tubes (Bertin) and 1 ml of RLT buffer (Qiagen) supplemented with 1%(v/v) Beta-mercaptoethanol. Tissue homogenate was passed through a QIAshredder homogenizer (Qiagen) and a volume that equated to 17.5 mg of tissue was extracted using the BioSprint™96 One-For-All vet kit (Qiagen) and Kingfisher Flex platform as per manufacturer’s instructions. Non-tissue samples were inactivated by placing samples into AVL buffer (Qiagen) and adding 100% ethanol. Extraction of these samples was performed using the BioSprint™96 One-For-All vet kit (Qiagen) and Kingfisher Flex platform as per manufacturer’s instructions.

Reverse transcription-quantitative polymerase chain reaction (RT-qPCR) was performed using TaqPath™ 1-Step RT-qPCR Master Mix, CG (Applied Biosystems™), 2019-nCoV CDC RUO Kit (Integrated DNA Technologies) and QuantStudio™ 7 Flex Real-Time PCR System (Applied Biosystems™). PCR amplicons were quantified against *in vitro* transcribed RNA N gene fragment standard. Positive samples detected below the lower limit of quantification (LLOQ) of 10 copies/μl were assigned the value of 5 copies/μl, undetected samples were assigned the value of 2.3 copies/μl, equivalent to the assays LLOD. For nasal swab, throat swab, BAL and blood samples extracted samples this equates to an LLOQ of 1.29 x 10^4^ copies/ml and LLOD of 2.96 x 10^3^ copies/ml. For tissue samples this equates to an LLOQ of 5.71 x 10^4^ copies/g and LLOD of 1.31 x 10^4^ copies/g.

Subgenomic RT-qPCR was performed on the QuantStudio™ 7 Flex Real-Time PCR System using TaqMan™ Fast Virus 1-Step Master Mix (Thermo Fisher Scientific) with forward primer, probe and reverse primer at a final concentration of 250 nM, 125 nM and 500 nM respectively. Sequences of the sgE primers and probe were: 2019-nCoV_sgE-forward, 5’ CGATCTCTTGTAGATCTGTTCTC 3’; 2019-nCoV_sgE-reverse, 5’ ATATTGCAGCAGTACGCACACA 3’; 2019-nCoV_sgE-probe, 5’ FAM-ACACTAGCCATCCTTACTGCGCTTCG-BHQ1 3’. Cycling conditions were 50°C for 10 minutes, 95°C for 2 minutes, followed by 45 cycles of 95°C for 10 seconds and 60°C for 30 seconds. RT-qPCR amplicons were quantified against an in vitro transcribed RNA standard of the full-length SARS-CoV-2 E ORF (accession number NC_045512.2) preceded by the UTR leader sequence and putative E gene transcription regulatory sequence. Positive samples detected below the lower limit of quantification (LLOQ) were assigned the value of 5 copies/μl, whilst undetected samples were assigned the value of ≤0.9 copies/μl, equivalent to the assays lower limit of detection (LLOD). For nasal swab, throat swab, BAL and blood samples extracted samples this equates to an LLOQ of 1.29 x 10^4^ copies/ml and LLOD of 1.16 x 10^3^ copies/ml.

For tissue samples this equates to an LLOQ of 5.71×10^4^ copies/g and LLOD of 5.14 x 10^3^ copies/g.

### Histopathology

Tissue samples from left cranial and caudal lung lobes, trachea, larynx, mediastinal lymph node, tonsil, heart, thymus, pancreas, spleen, liver, kidney, duodenum, colon, brain, vaccinating site (skin including subcutis and underlying muscle) and draining lymph node (left and right) were fixed in 10% neutral-buffered formalin and embedded into paraffin wax. 4 μm thick sections were cut and stained with haematoxylin and eosin (HE). Tissue slides were scanned and examined independently by two veterinary pathologists blinded to the treatment and group details.

For the lung, three sections from each left lung lobe were sampled from different locations: proximal, medial and distal to the primary lobar bronchus. A scoring system ^30^ was used to evaluate objectively the histopathological lesions observed in the lung tissue sections. The scores for each histopathological parameter were calculated as the average of the scores observed in the six lung tissue sections evaluated per animal.

Additionally, RNAscope in-situ hybridisation (ISH) technique was used to identify the SARS-CoV-2 virus in both lung lobes. Briefly, tissues were pre-treated with hydrogen peroxide for 10 mins (RT), target retrieval for 15 mins (98-101°C) and protease plus for 30 mins (40°C) (all Advanced Cell Diagnostics). A V-nCoV2019-S probe (Advanced Cell Diagnostics) targeting the S-protein gene was incubated on the tissues for 2 hours at 40°C. Amplification of the signal was carried out following the RNAscope protocol (RNAscope 2.5 HD Detection Reagent - Red) using the RNAscope 2.5 HD red kit (Advanced Cell Diagnostics). Appropriate controls were included in each ISH run. Digital image analysis was carried out with Nikon NIS-Ar software in order to calculate the total area of the lung section positive for viral RNA.

### Computed Tomography (CT) Radiology

CT scans were collected from sedated animals using a 16 slice Lightspeed CT scanner (General Electric Healthcare, Milwaukee, WI, USA) in the prone and supine position. All axial scans were performed at 120KVp, with Auto mA (ranging between 10 and 120) and were acquired using a small scan field of view. Rotation speed was 0.8s. Images were displayed as an 11cm field of view.

To facilitate full examination of the cardiac / pulmonary vasculature, lymph nodes and extrapulmonary tissues post-challenge, Niopam 300 (Bracco, Milan, Italy), a non-ionic, iodinated contrast medium, was administered intravenously (IV) at 2ml/kg body weight and scans collected immediately after injection and ninety seconds from the mid-point of injection. Scans were evaluated for the presence of COVID disease features: ground glass opacity (GGO), consolidation, crazy paving, nodules, peri-lobular consolidation; distribution - upper, middle, lower, central 2/3, peripheral, bronchocentric) and for pulmonary embolus. The extent of lung involvement was estimated (<25%, 25-50%, 51-75%, 76-100%) and quantified using a scoring system developed for COVID disease.

## Acknowledgements

Very special thanks to Amy Shurtleff for her help and support throughout this collaboration, Arun Kumar and Gerald Voss (CEPI) for their help and support and for critically reading the manuscript.

Thanks to Acuitas Therapeutics and Polymun Scientific for LNP formulations.

Thanks to Andreas Theß, Moritz Thran, Wolfgang Große for their support with generating mRNA constructs, Michael Firgens and Cibele Gaido for all their excellent work on project planning and communication and thanks to Igor Splawski for critically reading the manuscript. Thank you to Marilyn Aram, Rebecca Cobb, Breeze Cavell, Laura Hunter, Chelsea Kennard, Elizabeth J. Penn, Lauren Allen, Stephanie Leung, Emily Brunt, Stephen Thomas, Vanessa Lucas for assistance with sample analysis, and Susan Fotheringham, Nathan Wiblin and all Biological Investigations Group staff for animal husbandry and assistance with in vivo procedures. Our thanks also to Sara Wells and Claire Witham at MRC Centre for Macaques (A430).

## Author contributions

S.R., Y.H., S.O.M., M.J.D, S. S, M.C. and B.P. conceived and conceptualised the work and strategy. S.R., B.P., K.G., Y.H., F.V.G., H.E.H., S.L., D.N., J.P., K.A.R., R.J.W. and M.C., designed study setup, analysed and interpreted data. K.G. and D.H. supervised the study, F.J.S and E.L.R. performed histopathology analyses, S.R. wrote the manuscript. All authors supported the review of the manuscript.

---

**Supplementary Figure 1:**
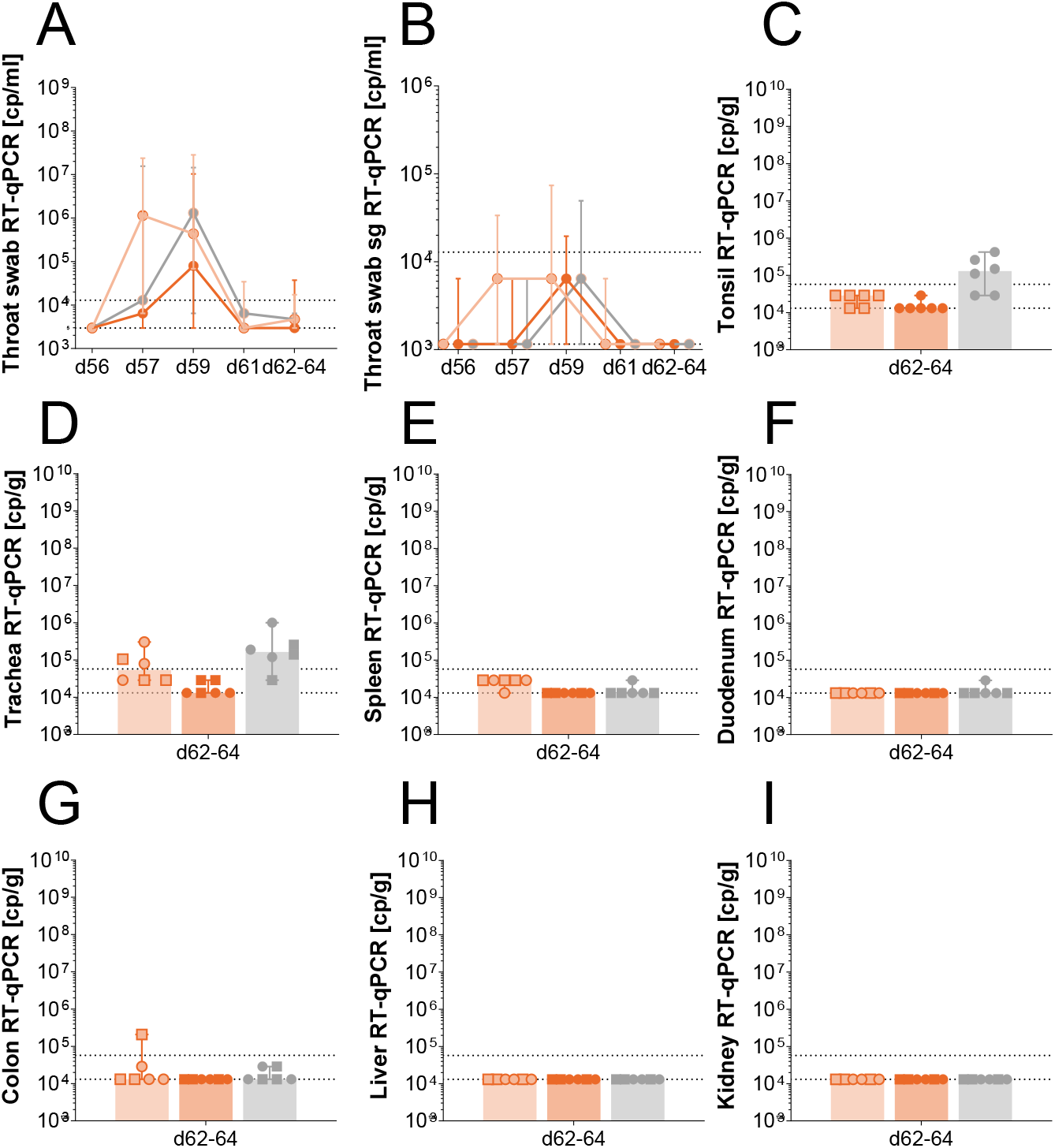
CVnCoV protects non-human primates from challenge infection (A) Throat swabs taken at different time points past challenge were analysed for copies of total viral RNA via RT-qPCR (B) Throat swabs taken at different time points past challenge analysed for copies of subgenomic RNA via RT-qPCR. Homogenised tissue derived from (C) tonsils (D) trachea (E) spleen (F) duodenum (G) colon (H) liver (I) kidney were analysed for copies of total viral RNA via RT-qPCR RT-qPCR Reverse transcription-quantitative polymerase chain reaction, sg subgenomic

